# Engineering *sHsp17* and *Hsp90* in *Zea mays* to Develop Thermotolerance

**DOI:** 10.1101/2025.11.26.690697

**Authors:** Usman Babar, Iqrar Ahmad Rana

## Abstract

**Background:** Genetically modified (GM) crops have shown a considerable success worldwide in improving farm incomes and reducing environmental pollution by limiting the use of synthetic chemicals. Maize is an important cereal crop of Pakistan and majorly grown in Punjab under Spring and Autumn seasons. There is a temperature shift in both seasons which bring a stress on crop plant, as a result of which the yield of the hybrids drops drastically due to failure of pollinations or heat coupled with drought causes reduced seed growth and hence less yield. This makes maize yield unstable over the years and keeps the farmer at risk. Current study was proposed keeping the above situation in mind. *shsp-17* from *Nicotiana tabacum* and *ATHSP-90* from *Arabidopsis thaliana* have been transformed in maize to play its role under biotic and abiotic stresses. Though maize plant also produces them, the target was to enhance the natural production of these proteins by genetic engineering in maize. *In planta* transformation protocols were equipped for the recovery of transgenic plants carrying reporter gene and gene of interest (*sHSP-17*). Putative transgenic plants for *HSP-90* were not achieved, at this we compared tissue culture/regeneration after bombarding immature zygotic embryos with selectable marker gene and found out that the said gene was perhaps deleterious for plant survival. The transgenic plants carrying reporter gene were checked for the presence of reporter protein by providing enzyme substrate which proved the transformation and expression of reporter gene for protein. The transgenic plants carrying *sHSP-17* were grown till maturity, confirmation for the presence of gene in maize genome was done using PCR and Southern blot while expression was checked using reverse transcriptase PCR. Seed of the transgenic plants were grown to get T1 generation which were found to be unable to attain maturity and produce progeny. It indicates that may be the expression of this gene too was deleterious for embryo survival and only those seeds could be developed where transgene was not available due to normal gene reshuffling during sexual reproduction.

## 1. Introduction

Maize is the fourth largest crop of the country after wheat, cotton and rice. It is grown as spring and autumn crop on an area of more than two million hectares with growing grain production rate of 5.39% on average per annum. It has increased from 705 thousand tons in 1971 to 7,800 thousand tons in 2020. According to USDA, the production has increased by 11% from last year due to increase in cultivation land but its production (per heactare) has dropped by 11% i.e., from 610 kg/hec to 544 kg/hec (USDA, 2021). In Pakistan, it is basically a food security crop with some industrial applications but worldwide, maize has a variety of uses including food, feed, and bioenergy. About 60 percent of the total maize production is used in poultry sector in Pakistan while only 25 percent is being used in agribusinesses. Rest of the crop production is used as fodder and feed for animals and humans (Rasheed, 2021).

In the life of every cereal, there comes a temperature shift between vegetative and reproductive growth stages. Abrupt changes in the climatic conditions during vegetative and reproductive growth stages affect plant performance negatively. These changes are quite frequent now under the climate change scenario. Being the C4 plant maize can efficiently use light energy and tolerate higher temperatures to some extent. April and September are critical months for spring and autumn maize crops respectively. Hot April and September coupled with drought stress severely affect grain yield. High temperatures result in pollen desiccation and reduced photosynthesis as the concerned enzymes like ROBISCO lose their activity severely at temperatures above 32.5 °C, similarly cell division of the endospermic tissues affects badly and results in the loss of grain weight to great extent (Aslam et al., 2013). Temperatures above 35 °C result in 101 Kg/ha/day reduction in grain yield (Saleem et al., 2013). In maize crop the yield is dependent upon the use of high yielding hybrids. The hybrids perform very well under suitable conditions but per hectare yield drops drastically the year temperatures go high especially at reproductive stage or if there is any kind of drought stress. Conventional breeding strategies have been employed to resolve the issue, but the success has not been great most of the time. This is because heat and drought stress tolerance is a complex phenomenon and is controlled by multigene families. With the emergence of the knowledge of biotechnology and genetic engineering it is possible now to scan many phenomena at the molecular level and then find the solutions. Many studies concerning the stress tolerance in general and “heat and drought tolerance” in particular, have shown an array of genes responsible. Heat shock proteins are a group of proteins which were identified at first in to play their role against heat stress (that’s why were given this name). Later on their function has been established more as general chaperons those bring rehabilitation in cell after stress. In this project we tried to express couple of genes expressing *hsp* proteins into maize by genetic transformation.

For efficient genetic transformation, reproducible regeneration systems are the prerequisite. In maize the regeneration studies started in the mid-eighties which culminated in the recovery of fertile transgenic in early nineties. The protocols were further improved through many efforts and model genotypes were identified for genetic transformation studies. A-188, B-173 and their hybrid Hi-II are the major genotypes which are targeted for maize transformations. Unfortunately, all of them grow in the temperate zone and tough to establish in tropical to subtropical climates of Pakistan. For maize genetic transformation little work is done in Pakistan and only a couple of studies from Centre of Excellence in Molecular Biology, Lahore exist. Both the studies were published in 2005 and 2006 respectively (Rafiq et al., 2005 and Rafiq et al., 2005 and 2006). In these studies, they tried to optimize the regeneration protocols and later transformed *Bt* gene using gene gun. In the present study, synthetic small heat shock protein (*Nt-hsp-17*) has been overexpressed in maize by *in planta* transformation and the protocols for regeneration and transformation in modern hybrids and open pollinated genotypes obtained from Maize and Millets Research Institute, Sahiwal has been well-established. These genotypes are otherwise high yielding but are susceptible against heat and drought stresses. On this *in planta* transformation protocols were tried which resulted in the development of transgenic plants with reporter genes as well as genes of interest.

## 2. Material & Methods

### 2.1. Genes Synthesis and Cloning

Gene sequences for *NT-sHSP-17* and *ATHSP-90* were synthesized by Genscript in cloning vector *PUC-57*-simple and confirmed by restriction digestion as well as PCR (Sambrook *et al*., 1989; Sambrook and Russel, 2001). These genes were cloned into binary vector *p7i-UG* using *Sac-I* and *Spe-I* restriction sites. The gene cassettes of *NT-sHSP-17* and *ATHSP-90* are shown in Figure 2.1.

**Figure 2.1.**
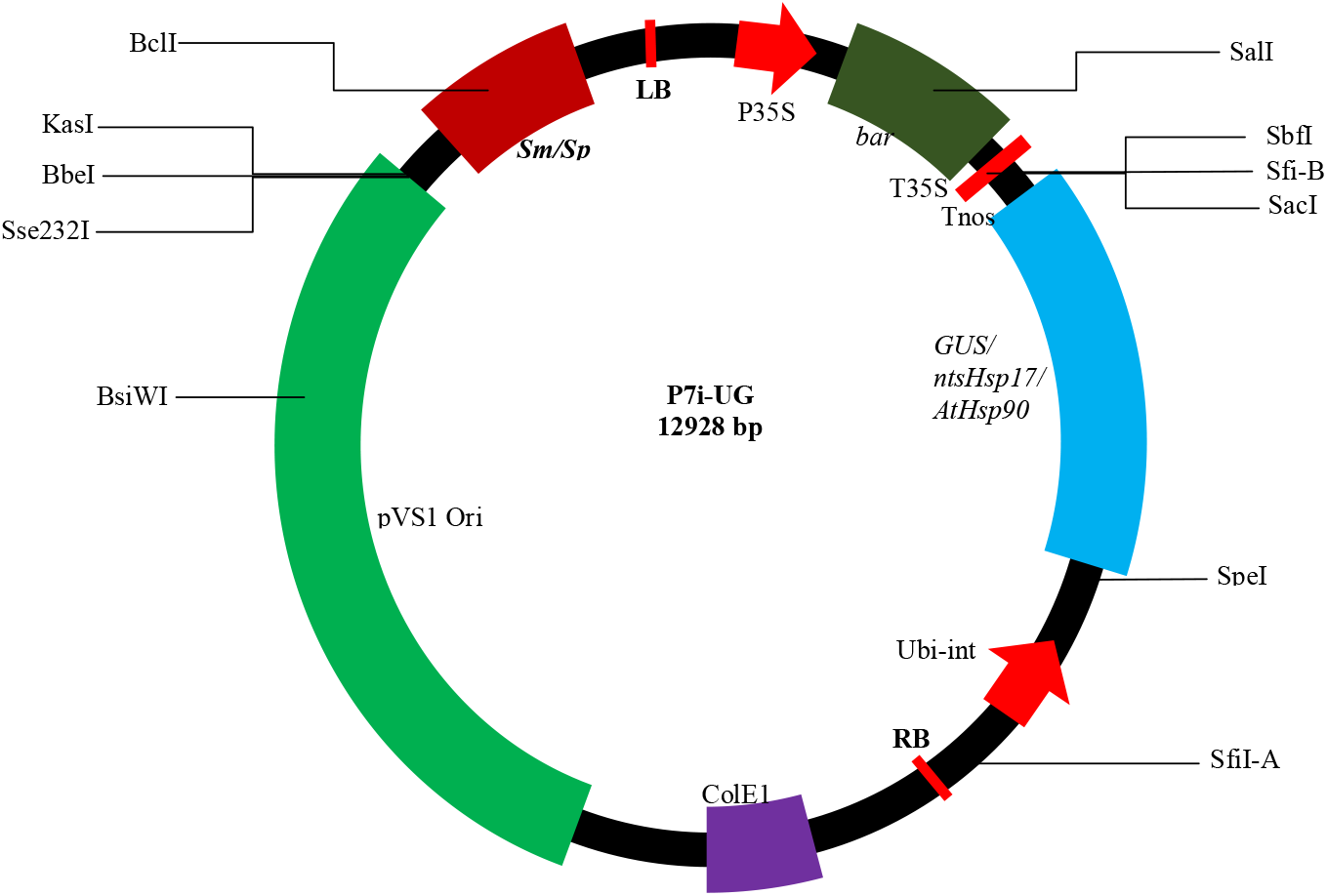
Gene Cassettes of *nt-sHSP-17* and *AtHSP-90*. As indicated in Figure 2.1, the genes of interest were cloned separately under ubiquitin promoter (*ubi*) and *nos* terminator, and the selectable marker gene *bar* was cloned under p35s and T35s.

**Figure 2.2.**
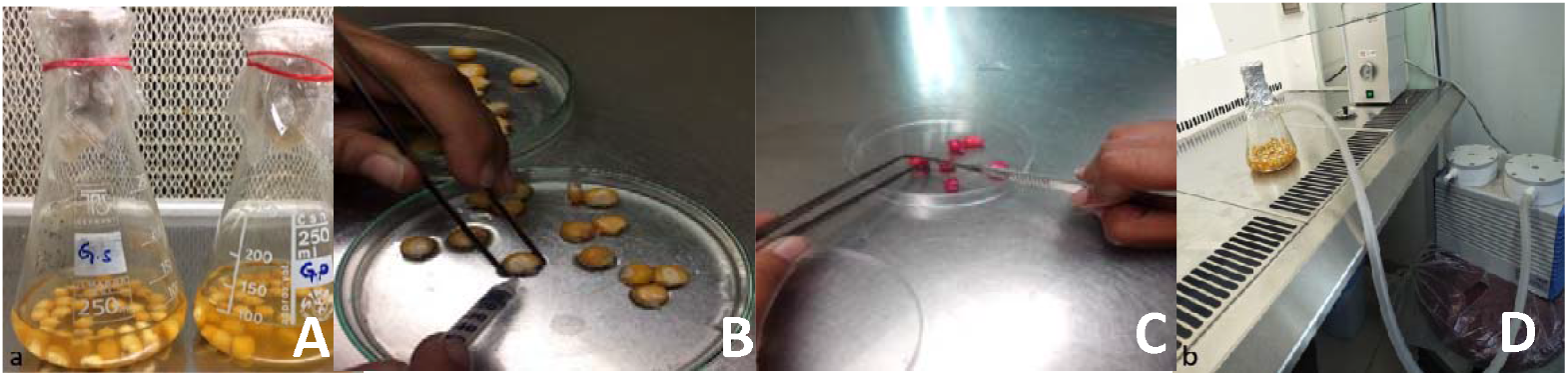
*In planta* Transformation Protocol by treating mature seeds: (A) Sterilization of maize seeds (B) Scarification of seeds (C) Seed puncturing (D) Co-cultivation of *Agrobacterium* inoculum via Vacuum infiltration

**Figure 2.3.**
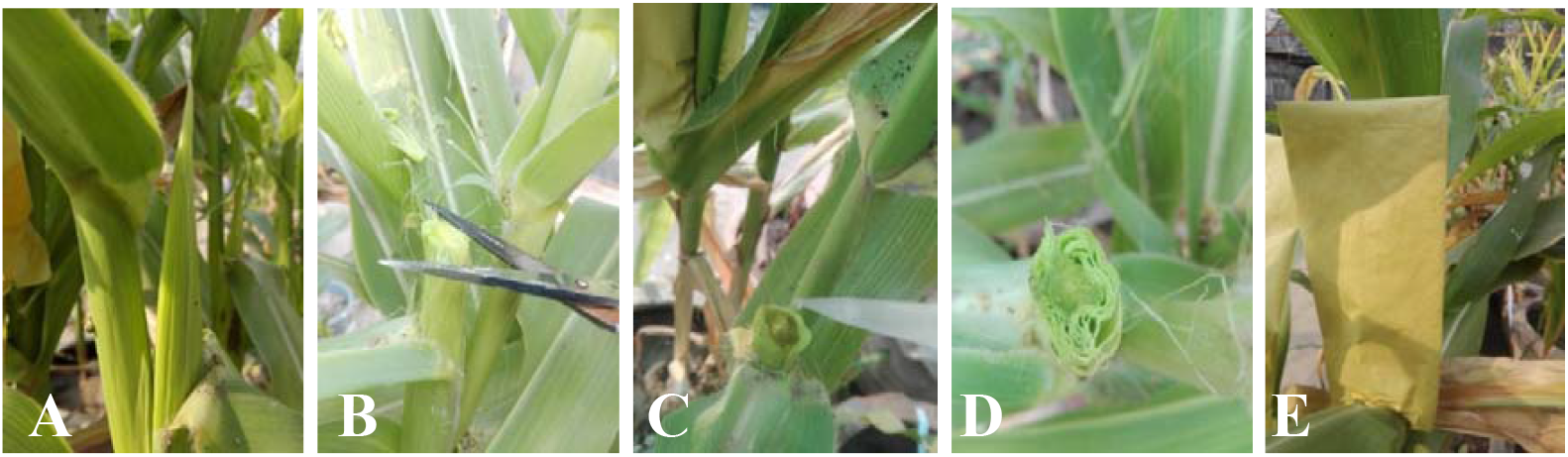
*In planta* Transformation Protocol: (A) Cob at initial silking stage (B) Cutting of apical region (C) Inoculation of Transformed *Agrobacterium* Culture (D) Artificial Pollination/Inoculation of transformed pollens (E) Covering of Cob

**Figure 2.4.**
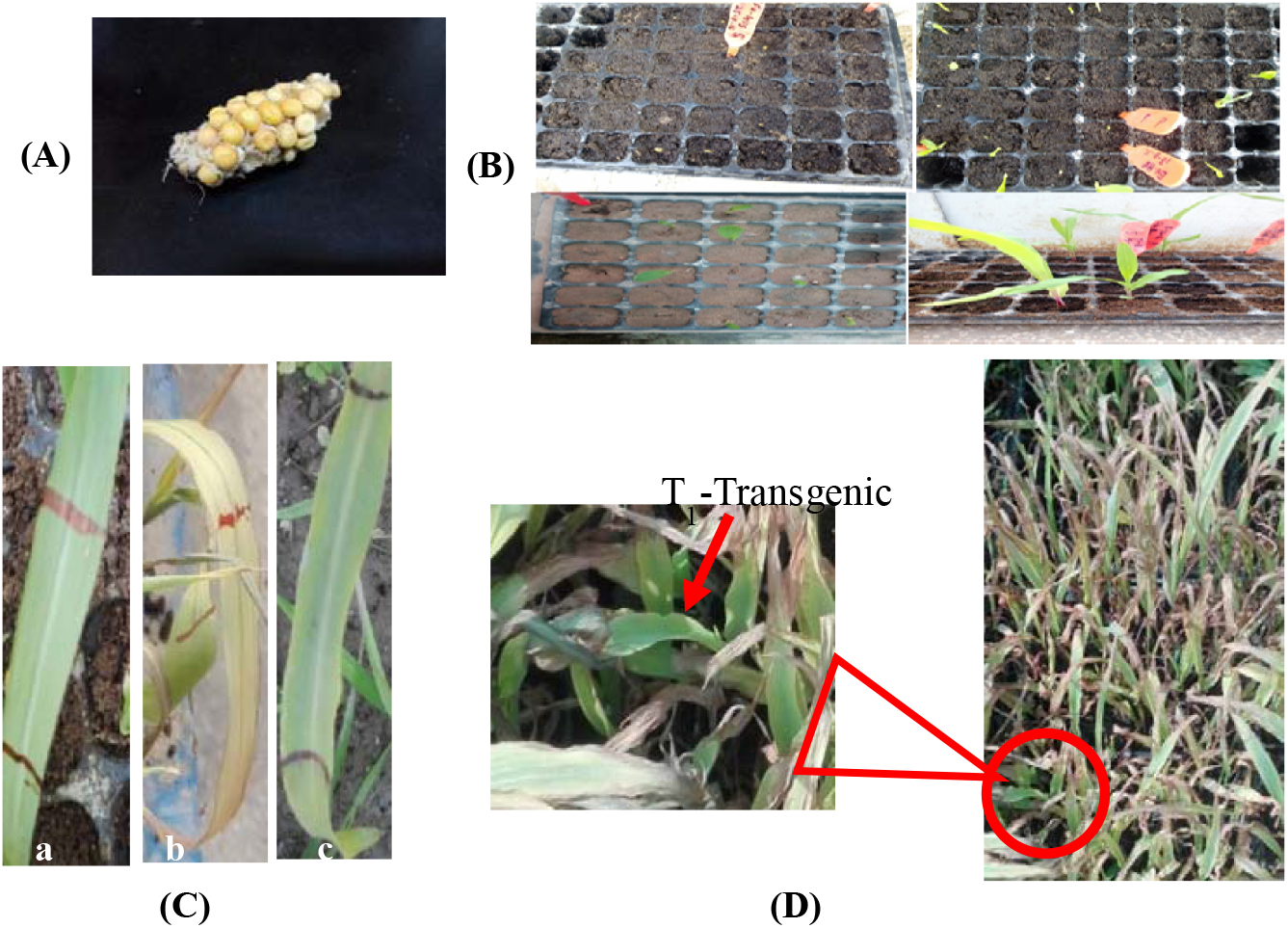
Screening of Putative Transgenics: (A) T_0_ artificially and selectively pollinated cobs (B) Sowing of T_0_ generation (C) Leaf-paint assay where, ‘a’ represents PPT-resistant plant, ‘b’ and ‘c’ represent PPT-susceptible plants, which refers to the transgenic and non-transgenics (D) Basta Spray on T_1_-generation

### 2.2. Plant Material

Three maize local genotypes viz EV-77, FH-949 and FH-1049 were used for *in planta* transformation. Quality seeds were collected from Maize and Millet Research Institute (MMRI), Sahiwal and Maize Research Station, Ayub Agriculture Research Institute (AARI), Faisalabad.

### 2.3. Agrobacterium-mediated Transformation Steps

Agrobacterium mediated transformation was optimized following majorly Frame et al., 2002 and Vega et al., 2009.

### 2.4. *In planta* Transformation by Scarification and Puncturing Method

### 2.5. *In planta* Transformation through Pollen Tube Method

### 2.7. Transformation Trials

According to protocol mentioned in heading 2.5, in both treatments, total 960 seed were sown. Out of them 480 seeds were applied first treatment named scarifying. Other 480 seeds were applied second treatment named puncturing.

According to protocol mentioned in heading 2.6, 90 cobs were selected for conducting the experiment 30 were treated with induction media-I (IM-I) (Hensel *et al*., 2009), 30 were treated with induction media-II (IM-II) (Patel, 2007) while rest of the 30 cobs were treated with induction media-III (IM-III) (Das and Joshi, 2011). The recipes of induction media are mentioned in Table 2.2.

**Table. 2.1.** Induction Media Preparation Recipe.

| Induction Media I (IM-I) | Induction Media II (IM-II) | Induction Media III (IM-III) |
| --- | --- | --- |
| 1.1 g N6 medium with vitamins<br>27.4 g Sucrose<br>14.4 g Glucose<br>0.2 g L-Proline<br>0.2 g MES<br>0.6 mg 2,4-D<br>0.05 mg Silver Nitrate<br>5 μL Acetosyringone<br>100 mL Autoclaved DH<sub>2</sub>O<br>pH: 5.2 | 1.1 gN6 medium with vitamins<br>0.01 g Myo-inositol<br>0.01 g Casein hydrolysate<br>6.84 g Sucrose<br>3.6 g Glucose<br>0.28 g L-Proline<br>0.74 g Fe-Na-EDTA<br>0.8 mg 2,4-D<br>5 μL Acetosyringone<br>100 mL Autoclaved DH<sub>2</sub>O<br>pH: 5.8 | 0.5 mg MS-media with vitamins<br>5% Sucrose<br>0.075% Tween 20<br>5 μL Acetosyringone<br>100 mL Autoclaved DH<sub>2</sub>O<br>pH: 5.7 |

\*Concentration of Acetosyringone (AS) used in all the media was 100 $\mu$ mol L<sup>-1</sup>.

### 2.8. Screening of Putative Transgenics

All the T_0_ seeds from both transformation trials were sown into the compost-containing trays under controlled conditions and screening process was done in two ways under 200 mgL^−1^ basta or Phosphinothricine (PPT) selection pressure i.e., leaf-paint assay and spray. The PPT herbicide application was repeated for two to three times in order to avoid the ambiguity (Rajasekaran *et al*., 2017). The seeds obtained from T_0_ generation were sown again and T_1_ was screened for transgenics as well.

## 3. Results

### 3.1. Molecular Characterization

PCR analysis of 38 suspected plant DNA samples was performed and the amplification activity was found in 5 plant DNA samples. Where, Figure 3.1 shows the molecular screening of transformants among suspected plants along with the PCR confirmation of integrated *sHSP-17* gene within maize genome of these plants.

**Figure 3.1.**
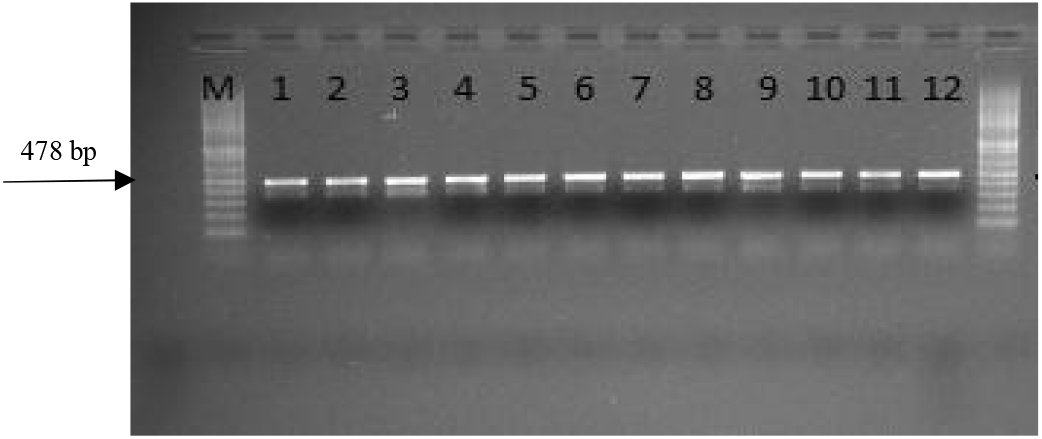
PCR results of PPT-resistant plants. Lane designation: M: 1 Kb Ladder. (1-12) candidate samples. (+) Positive control from *Hsp 17*. -ve is PCR run on DNA isolated from non-transformed maize plant.

### 3.2. Southern Blot Chromogenic Detection

The gene-specific probes designed were found to be hybridized with PCR+ DNA samples. Chromogenic detection was made with the action of NBT/BCIP. Figure 3.2 shows the chromogenic detection of probe-to-DNA-specific hybridization.

**Figure 3.2.**
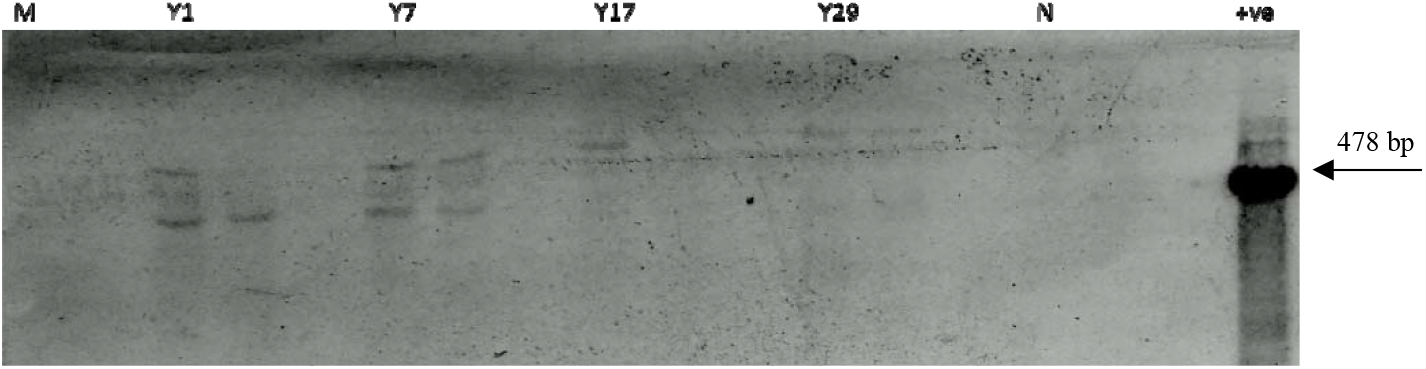
Chromogenic Detection of probe-to-DNA-specific Hybridization. Y1, Y7, Y11 and Y17 are the DNA samples of PCR+ candidate plants, N=non-transformed plant samples, P=p7i-*sHSP*-UG Plasmid and M= 100 bp Marker.

### 3.2. Expression Analysis

For the expression analysis, gene specific primers were used, Figure 3.3 shows the presence gene specific amplification of the same size in 4 tested samples as shown by the plasmid carrying the gene of interest. There was no amplification in negative control which excludes any contamination or amplification of endogenous fragment.

**Figure 3.3.**
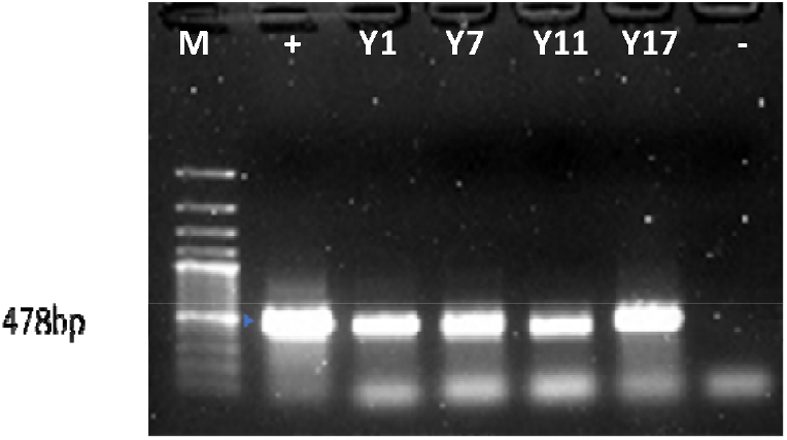
Amplification of *shsp-17* gene from transformed plants cDNA. M=1Kb DNA marker, += fragment amplified from plasmid, 1, 4, 5, and 6 = fragments amplified from cDNA synthesized from transgenic plants verified through selection pressure, PCR and Southern Blot analysis.

### 3.3. Gus-Assay

In order to check if the selectable marker gene went silent plants selected for p7i-UG carrying *bar* and *gus* gene were selected through basta application and gus assay was performed in comparison with non-transformed plant tissue. Fig 3.4 represents the pollens harboring *uidA* gene which is responsible for staining of transformed plants. As shown in Figure 3.5, *uidA* carrying plant tissues showed development of blue color while non-transformed plant tissues showed no such color development. The reason of non-selectivity may be the unavailability of *Nt-sHSP-17/bar* gene in T_1_ generation possibly due to toxicity of *Nt-sHSP-17* gene product in embryo development.

**Figure 3.4.**
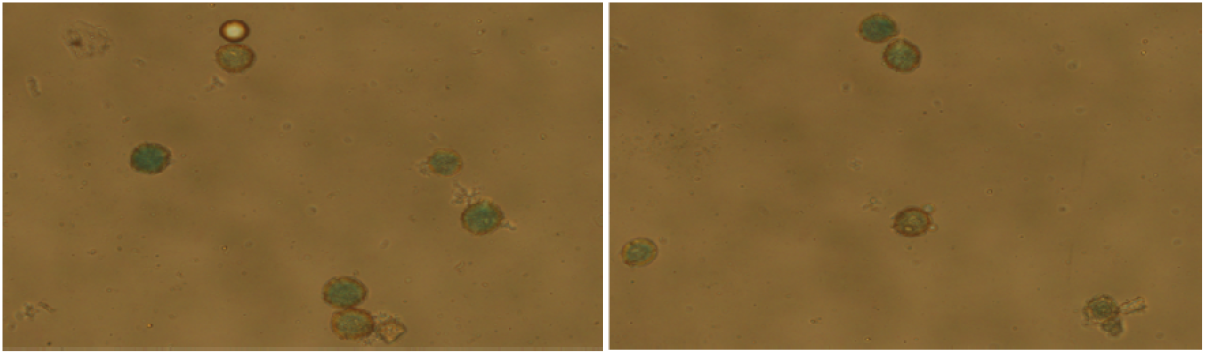
*Gus* staining of pollens transformed with uidA gene

**Figure 3.5.**
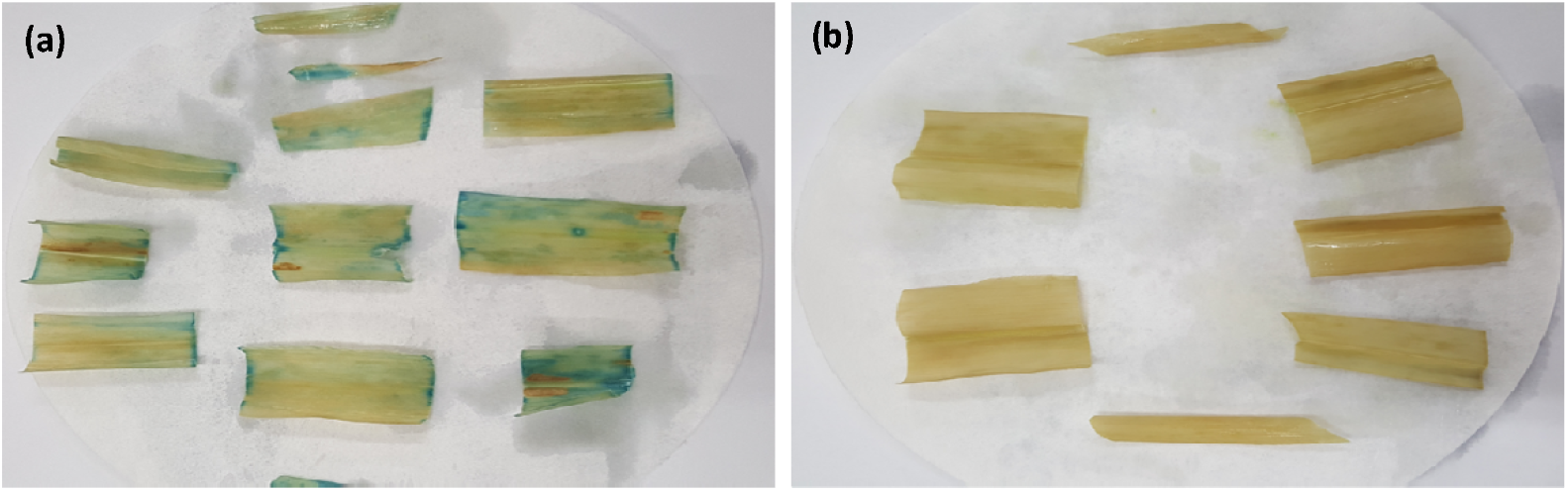
*gus* expression analysis of *uidA* positive transgenics from T_1_ generation and non-transgenic plant leaves. (a) survived plantlets (b) damaged plantlets

**Figure 3.6.**
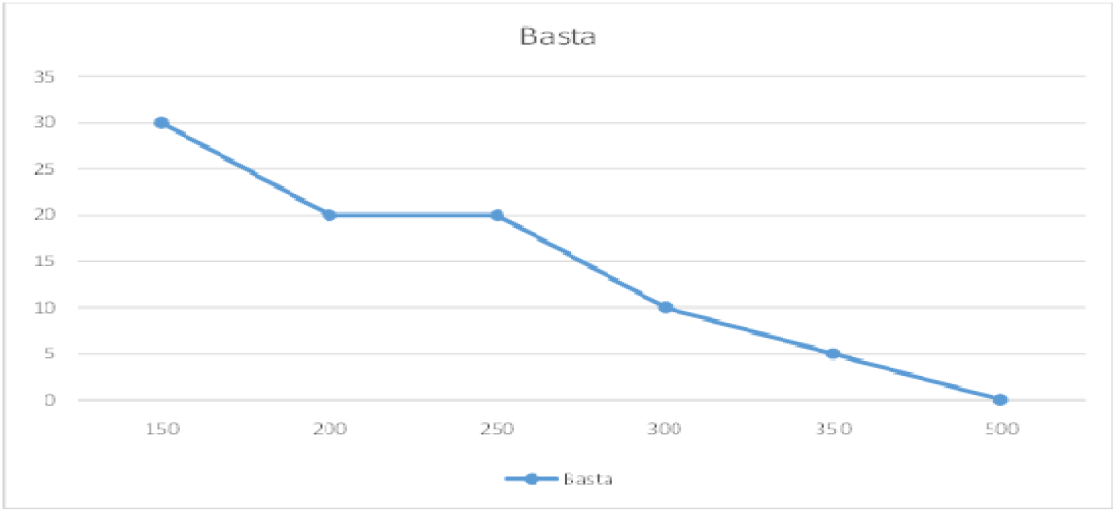
Kill curve analysis showing death percentage versus basta (ppt) concentration in mg/l

### 3.4. Screening of putative transgenic plants by BASTA application

Before going for screening of transformed material non transformed plants were sprayed with various concentrations of phosphinthricin (ppt) and it was noticed that total plant death occurs at 500 mg/l ppt but plants generally withers at 150-200 mg/l in a 7-10 days. When ppt is applied on a localized area with cotton swab local tissue starts bleaching even at 100mg/l ppt in 7-10 days post inoculation. Graph below shows the kill curve analysis.

In Trial II, The AGL-treated germplasm (90 cobs) were collected and the obtained seeds (247) of all the induction media were then sown and grown to maturity for phenotypic screening of putative transformants that was done under Basta (Glufosinate ammonium) selection pressure (200 mg L^−1^). Only the successful transformants with expressed *bar* gene have showed tolerance against this herbicide. 38 plants were phenotypically screened out of the 247 plants. 13 Basta-resistant plants while 25 partially or semi-resistant plants were obtained.

### 3.5. Determination of Transformation Efficiency

#### 3.5.1. Puncturing and scarifying the maize seeds

In Trial I, out of 960 seeds, 206 plantlets appeared 95 from scarified seeds and 111 from punctured seed. Development proportions were 19.7% (95/480) and 23.1% (111/480) for scarifying and the puncturing-treated seeds, respectively. At three to five leaf stage, basta was applied two times for the selection purposes and 28 plantlets survived (12 by scarifying treatment and 16 by puncturing treatment. The data is shown in table 3.1.

**Table. 3.1.** Transformation experiment Data at T_0_ generation.

| Treatment Type | No. of treated growing seeds | Germination level | PPT-tolerance | PCR Positive Plants |
| --- | --- | --- | --- | --- |
| Scarifying | 480 | 95 (19.7%) | 12 (12.6%) | 2 (16.6%) |
| Puncture | 480 | 111(23.1%) | 16 (14.4%) | 4 (25.0%) |
| Total | 960 | 206 (21.4%) | 28 (13.6%) | 6 (21.4%) |

**Table. 3.3.** Comparison of Transformation Frequency on basis of Induction Media.

| M | Treated Cobs | Obtained Seeds | Screened Plants | Transformation Frequency |
| --- | --- | --- | --- | --- |
| IM-I | 30 | 92 | 4 | 4.3% |
| IM-II | 30 | 43 | 1 | 2.3% |
| IM-III | 30 | 112 | 1 | 0.8% |
| Avg. |  |  |  | 2.46% |

Seed germination reduced drastically in the trays as shown above in Figure 2.4 and also evident from table 3.1. In total 21.4% seeds could grow from a total of 960 treated seeds (19.7% scarified and 23.1% punctured seeds could grow). The obtained transgenic were from EV-77 and FH-1049.

### 3.5.2. Transformation through Induction Media and Comparison of their Efficiency

In Trial II, the transformation of plants was carried out by applying induced Agrobacterium in induction media to obtain enhanced transformation efficiency. 30 plants were treated with each induction media and the seeds obtained from them were taken as subject for the determination of transformation frequency and efficiency. The applied transformation protocol was quite successful and the transformation frequency of 2.46% was achieved as a result of this research experiment. The explants treated with induction media I (IM-I) showed 4.3% transformation efficiency, plants treated with induction media II (IM-II) showed 2.3% transformation efficiency while induction media III (IM-III) showed 0.8% transformation efficiency. Hence, it is clearly stated that IM-I performed best in transformation process, it plays a vital role in the enhancement and effectiveness of the technique. Similar to trial I, obtained transformants were from EV-77 and FH-1049, however no transformant was obtained from FH-949 genotype.

In Trial I, puncturing method was observed to be performing way better than the scarification likewise, IM-I produced the most promising results in Trial II. However, the transformation efficiency determined in both experiments has revealed that the transformation trial II is most likely for undergoing transformation in the maize as its transformation frequency as well as efficiency was observed to be greater than that of trial I.

### 3.6. T_1_ generation of the Transgenic plants and screening of transgenics

Through *in planta* transformation plants expressing *Nt-sHSP-17* and *uidA* gene for *gus* expression were achieved and extended to next generation after molecular analysis. As shown in Figure 3.7 seed of each plant was sown and sprayed with basta in order to see gene reshuffling in next generation.

**Figure 3.7.**
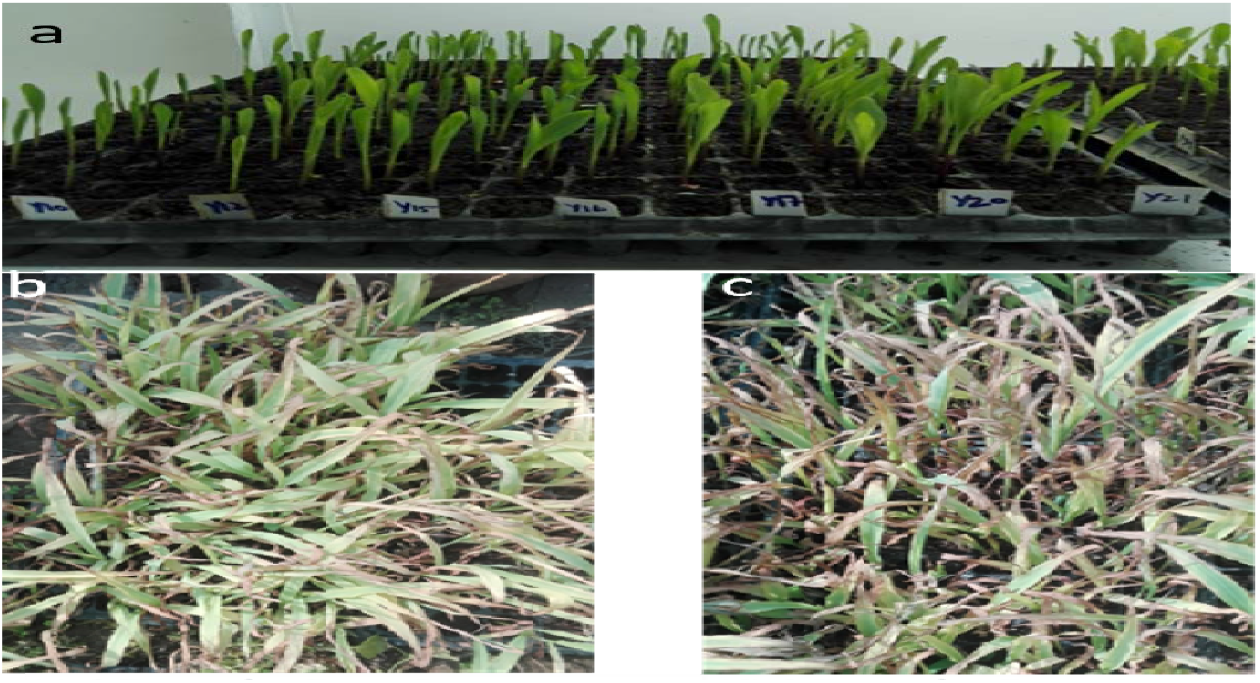
Treating T_1_ generation of transgenic plants with selection pressure (basta). (a) just after germination. (b,c) 7, 10 days after spraying basta.

As shown in Figure 3.7, it was observed that all the plants overexpressing *shsp17* could not be selected in next generation either due to reshuffling of said gene construct, and/or silencing of selectable marker gene bar. Only 3-5 plants were selected in T_1_ generation but these plants too were showing abnormal growth and could not reach maturity and bolted before that and no seeds were developed from them.

## 4. Discussion

Agrobacterium mediated *in planta* transformation was already reported in many crop plants including maize. Majorly two methods were identified to be successful in maize in the recently reported survey. Abhishak *et al*., 2016 reported the recovery of transgenic seed by developing chimeric transgenic plants. The did it by injuring the germinating maize seeds by puncturing and scarification. They expected the development of chimeric transgenic tissues if such seeds are treated with agrobacterium. The seeds if developed from such chimera will be transgenic seeds. Working on this hypothesis they recovered transgenic plants in generations of injured agrobacterium treated seeds. In another experiment Chumakov *et al*., 2006 reported the pollen tube mediated pathway by treating young silks with agrobacterium and immediately pollinating them. The hypothesis they worked on was either pollens will get transformed or agrobacterium will slide to egg cells and transform it. In either case the developed embryo will be transgenic. By using this approach, they recovered transgenic plants. These results were reconfirmed by Mamontova *et al*., 2010.

The *gus* gene was cloned in p7i-UG by using Sac-I and Spe-I restriction sites. It was planned to use these sites, to replace *gus* with genes of interest. Therefore, *Spe*-I and *Sac*-I were kept as additional sites, on 5’ and 3’ end of our genes of interest as additional sequences when getting these genes synthesized from Genscript USA. This kind of replacement strategy is already used by many in previous studies and it works very well (Rana *et al*., 2012, Oldach *et al*., 2002). It was done by using the recommended protocols and the integration of genes of interest (*Nt-shsp-17* and *At-hsp-90*) into expression vectors was also confirmed. The confirmation was done both by restriction digestion, sequencing, and PCR analysis.

*In planta* transformation was tried using two methodologies, the first method involved the development of transformed chimeras in the tissues of germinating embryos by agrobacterial vacuum infiltration. The other *in planta* strategy was based on pollen tube mediated transformation. Using both the strategies transgenic plants expressing *Nt-sHSP-17* and *UidA* gene were achieved. The earlier strategy where chimeras were developed, was based on reports by Wang *et al*., 2007 and Abhishek *et al*., 2016. They co-cultivated injured seed embryos with agrobacterium. To do better interaction with agrobacterium vacuum infiltration which has been reported successfully in Arabidopsis by many authors (Ye *et al*., 1999; Desfeux *et al*., 2000) was tried. In *brassica rapa* floral dip and vacuum infiltration were tried to achieve an overall transformation frequency of 0.15% while vacuum infiltration resulted in doubling of the transformation frequency to 0.3% (Hu *et al*., 2019). In our case when simple cocultivation of was tried there was almost no reduction in germination of treated seeds and chimeras were obtained only in few plants that could not be traced in next generations. On vacuum infiltration there was no germination upon 10 minutes of vacuum infiltration to germinating seeds while reduced germination was observed when seeds were infiltrated for 5 minutes on vacuum. In the next generation the plants showing chimeras were screened and at least three plants with abnormal growth were screened positive. These plants did not reach maturity.

The second *in planta* transformation strategy involved pollen/pollen tube mediated transformation. Maize pollens proved to be readily transformable in this research hence they were cocultivated on the silks with the idea that either they get transformed and fertilize the eggs later or carry agrobacterium to female parts as through published literature it is known that agrobacterium targets female parts during floral dip in model plant Arabidopsis (Desfeux *et al*., 2000). These protocols were modified from Mamontova *et al*., 2010 and Chumakov et al., 2006. Activated agrobacterial cells were inoculated onto receptive silks in three different induction media types. These media are more plant friendly, and out of the three media IM-I performed better (10% transformation efficiency) which contained N6 salts, sucrose (27.5g/l), maltose (14.4g/l), Acetosyringone (100mM), 2,4-D (2mg/l) and Silver Nitrate compared to others who either lack these nutrients or had them in low concentrations (Explained in Methodology). These media have been used by many for inoculating agrobacterium to plant tissue increase interaction and later the transformation efficiency. Using this methodology at least 15 plants were seen as PCR positive after selectable marker-based screening. Out of these 15 plants 5 plants were checked by Southern blot analysis for identification of copy number, which seemed single in all cases. 5 out of 15 were with abnormal growth and bolted very early with infertile pollens and silk less cobs. The others developed cobs and reduced number of seeds which were screened in next generation and mostly could not survive against selection pressure. This was possibly due to the fact that transformed embryos could not survive or were abnormal in growth and did not reach maturity, and only non-transformed developed seeds died against selection pressure in next generation.

Current model of sHSP chaperons suggest that they assemble into large homo-oligomer and bind to denatured proteins in ATP independent manner then *HSP-70* and *HSP-90* cooperate with the complex in ATP dependent manner to convert them into functional form (Xu *et al*., 2011). The purpose of conducting this research was to separately transform *sHSP-17* and *HSP-90* into maize and then bring them together through crossing expecting a genotype highly affecting in rescuing proteins degraded by heat/drought stress. The literature supported this strategies and increased resistance by transforming individual *HSP* genes are reported in many plants like rice, barley, and tobacco etc. (Murakami *et al*., 2004; Poonia *et al*., 2020; Jiang *et al*., 2020). In our case *HSP-90* construct looked toxic and *sHSP17* gene got overexpressed, but abnormal phenotype was shown by some plants in T_0_ while others were normal. In T_1_ generation, only a few plants survived and those too were with abnormal phenotype and could not reach maturity. Hence the impact of transgene on stress tolerance could not be studied. There could be a few possible reasons of finding abnormal plants in T_0_ and T1 generations. The constitutive expression of *Nt-sHSP-17* in maize somehow interferes with reproductive system, fertilization, and embryo development. Constitutive expression of *sHSP-17* gene may also have interfered with certain metabolic pathways resulting in growth inhibition and death of developing embryos. The literature reports the differential expression of certain heat shock proteins during reproductive organs development like in fertilization and pollen and egg development in plants (Saeed *et al*., 2019). On the other hand, in overexpression of heat shock protein in Drosophila and mice cells resulted failure of fertilization and embryo development (Neuer *et al*., 2000; Silbermann and Tater, 2000).

## Conclusion

It is concluded from the said study that for efficient transformation and trait development through genetic engineering the identification of model genotypes in local tropical maize is of utmost importance. Secondly, the identification of inducible/tissue specific promoters can also improve the success of transformation experiments in crop plants. This project was hypothesized upon the introduction of heat shock protein genes (*Nt-sHSP-17* and *At-hsp-90*) into model maize genotypes, the model genotypes were exported but could not be maintained locally because they were bred and adapted to tropical environment only. Hence, one option could be the use of controlled greenhouses which were not currently available at UAF. Genes got synthesized, cloned into expression vectors. Extensive regeneration studies, transformation parameters optimizations were undertaken, transformation through agrobacterium was tried but no transgenic plant was recovered from T_1_ generation. Transformation experiments were done using mature and immature embryos as explants. Later on *in planta* transformations were optimized using a couple of methodologies (Induction of chimera, and pollen tube mediated transformation). *In planta* transformation proved to be successful both with Gene of interest (*Nt-sHSP-17*), optimization construct carrying *bar* and *gus* genes. *Gus* genes were successfully screened in next generation while gene of interest plants occasionally showed abnormal phenotypes in T_0_ generation and could not survive in T_1_ generation. From here we concluded that constitutive expression of said protein in maize plant interferes with certain essential metabolic pathway which resulted in occasional abnormal phenotypes even in T_0_ plants and in T_1_ none of the plants with homozygous or hemizygous state could survive and only the seeds with reshuffled transgenes survived.

The development on gene synthesis and cloning into expression vectors was done successfully. From the above discussions and the current literature reviewed it is evident that maize can be efficiently transformed after optimizing the basic parameters of transformation. The idea of mature seeds or taking mature embryos from the dried seeds and using them for transformation studies is not supported by the literature.

Worldwide; model maize genotypes (HI-II hybrid and its Inbred A-188 and B073) for transformation have been bred and they grow better under temperate climate only. Such genotypes should also be identified from tropical germplasm, their regeneration and transformation protocols should be finetuned further. The transformation protocols tried in the current attempt may work as baseline for the identification of such germplasm/genotypes. This will ensure the success of transformation experiments in those areas of the world where only tropical germplasm is cultivated. Same principle is followed in China for transformation of cotton, wheat, maize and other species. In addition, Heat shock proteins should be expressed under inducible promoter only. This will reduce the risk of interference of such proteins with endogenous proteins if expressed under constitutive promoter. The work is already under process for cloning of these genes under soybean heat inducible promoter.

